# Deep graph convolutional neural network for one-dimensional hepatic vascular haemodynamic prediction

**DOI:** 10.1101/2024.08.13.607720

**Authors:** Weiqng Zhang, Shuaifeng Shi, Quan Qi

## Abstract

Hepatic vascular hemodynamics is an important reference indicator in the diagnosis and treatment of hepatic diseases. However, Method based on Computational Fluid Dynamics(CFD) are difficult to promote in clinical applications due to their computational complexity. To this end, this study proposed a deep graph neural network model to simulate the one-dimensional hemodynamic results of hepatic vessels. By connecting residuals between edges and nodes, this framework effectively enhances network prediction accuracy and efficiently avoids over-smoothing phenomena. The graph structure constructed from the centerline and boundary conditions of the hepatic vasculature can serve as the network input, yielding velocity and pressure information corresponding to the centerline. Experimental results indicate that our proposed method achieves higher accuracy on a hepatic vasculature dataset with significant individual variations and can be extended to applications involving other blood vessels. Following training, errors in both the velocity and pressure fields are maintained below 1.5%. The trained network model can be easily deployed on low-performance devices and, compared to CFD-based methods, can output velocity and pressure along the hepatic vessel centerline at a speed three orders of magnitude faster.

**Author summary:** When using deep learning methods for hemodynamic analysis, simple point cloud data cannot express the real geometric structure of the blood vessels, and it is necessary for the network to have additional geometric information extraction capability. In this paper, we use graph structure to express the structure of hepatic blood vessels, and deep graph neural network to predict the corresponding hemodynamic parameters. The graph structure can effectively express the geometric information of hepatic blood vessels and the topology of branch blood vessels, which can effectively improve the prediction accuracy with strong geometric generalisation ability. The results show that the method achieves the highest prediction accuracy in the one-dimensional hepatic vessel blood flow simulation dataset, and the experimental results on the human aorta also show that our method can be effectively applied to the blood flow simulation of other vascular organs.

## Introduction

Hepatocellular carcinoma (HCC) is the most common primary hepatic malignancy and the leading cause of cancer-related deaths worldwide [1], and hemodynamic calculations of the hepatic vasculature play an important role in both the treatment and postoperative period of HCC [2–4]. one of the ways to obtain it is to utilize the CFD method [5, 6], after the calculations the hemodynamic parameters inside the vessel can be obtained, however, the CFD method requires a large amount of computational resources, which limits its application in clinical practice.

Deep learning is an alternative to CFD, where trained network models allow for rapid hemodynamic parameter prediction [7–9]. Li et al. [10] used neural networks to predict the haemodynamics of different cerebral aneurysms before and after blood flow steering stent implantation to provide guidance for interventional procedures for cerebral aneurysms. Wang et al. [11]predicted arterial blood flow in the foot based on plantar skin temperature mapping, and altered blood flow in the foot is an important indicator of early diabetic foot complications. Yuan et al. [12] used a convolutional neural network to predict drug trajectories and concentration fields in chemoembolisation treatments to aid in directing drugs to tumour sites. Raissi et al. [13, 14] proposed physics-informed neural networks (PINNs), which add physical information for constraints to the data-driven neural networks, enhancing the interpretability of the neural network. However, PINN is not flexible enough in defining the domain to satisfy the diverse geometric features of blood vessels, and the anatomical shapes of patients’ liver blood vessels usually have large geometrical variability due to different medical conditions, and neural network models based on 3D blood flow simulation data are also highly dependent on the arithmetic power of the device due to the size of the data.

One solution is to use data-based reduced-order models (ROMs) [17–19]. Reduced order modeling is a technique to transform a high-dimensional problem into a low-dimensional problem, which reduces the dimensionality of the simulation by retaining the main flow characteristics and key information.The use of ROMs reduces the dependence on high-performance equipment in the training process by reducing the size of the data. Drakoula et al. [20] used singular value decomposition to obtain a high-fidelity solution for 3D hemodynamic parameters, and the results demonstrate the predictive ability of the method for hemodynamics as well as the training efficiency. siena et al. [21] reduces the dimensionality of the data by means of the Principal Orthogonal Decomposition method, and the training of the neural network by means of the reduced dimensionality of the data can greatly shorten the time required for the training. It is possible to quickly and reliably numerically simulate the blood flow patterns occurring in a patient-specific coronary artery system. fresca et al. [22] utilized the POD method to provide accurate predictions of blood flow patterns in cerebral aneurysms in real time. Data-based downscaling models are more suitable for data with some structure or repetitive patterns. For highly nonlinear or complex data sets, as with PINNs, it is sensitive to changes in geometry.

Another solution is geometry-based ROMs, where neural networks are trained using one-dimensional hemodynamic simulation data from blood vessels [23–25],The trained network can predict hemodynamic parameters along the centerline of the patient’s vascular anatomy model. In CFD, the Reynolds number (Reynolds number) is the ratio of inertial force to viscous force of a fluid, and when the Reynolds number is small, the viscous force is small for the perturbation of the flow velocity in the flow field, and vice versa.Xiao et al. [26] A comparison was done for the 3D and 1D problem, and the difference between the 1D and 3D carotid arteries (Reynolds number of 748) was smaller than the difference between the aorta (Reynolds number of 7140), and the authors addressed this problem by reducing the inflow into the aorta to match the Reynolds number of the baseline carotid model, and found that the average error was reduced, and in vascular systems with smaller Reynolds numbers, the error between the 1D hemodynamic model and the 3D is also the smaller, i.e., blood flow is predominantly advective.Pfaller et [27] compared the results of one-dimensional simulations with those of three-dimensional simulations, and observed an average relative error of 1.8% for pressure and 3.4% for flow velocity. In the experiments of Kennedy et. [28] a Reynolds number of 1,150 in the hepatic vasculature was derived as valid, and other studies have demonstrated that the maximum Reynolds number in the hepatic vasculature is not more than 1800 [29]. Although one-dimensional models are not a true substitute for 3D models, they are easy to implement, time-saving, less demanding on computer hardware, and capture the main features observed in 3D and experimental models [30–32]. We can conclude that one-dimensional hemodynamic simulations targeting the hepatic vasculature yield similar results to three-dimensional simulations and that the scheme is feasible.

For numerous AI for Science applications, geometric graphs are a powerful and versatile representation that can be used to represent a multitude of physical systems, including small molecules, proteins, crystals, and physical point clouds. For blood vessels, the physical system is the same no matter how the vessels are moved or rotated in space. Compared to traditional neural networks trained from discrete point clouds that need spatial location information to extract geometric features, graph neural networks are more concerned with the geometric links of the blood vessels and the interactions between the nodes, and the trained graph neural networks have higher accuracy and better geometric generalization performance. For example, Suk [33, 34] et al. used a graph neural network to predict the velocity and directional wall shear stresses at the vertices of a vascular mesh and achieved more accurate and efficient results than PointNet++. Pegolotti et al. [35] represented the centerline of the blood vessel as a graph structure, and predicted the velocity and pressure at the graph nodes by a graph neural network, maintaining an accuracy of over 98% and 97% for pressure and velocity.

In this paper, we use a deep graph neural network for the prediction of one-dimensional hepatic vessel hemodynamics, constructing the centerline model of a complex hepatic vessel as a graph structure, with the points on the centerline as the graph nodes and containing the corresponding features, embedding the boundary conditions into the features of the graph nodes, and using the real geometric attributes between the nodes as the edge features in the graph. In this paper, based on the deep graph neural network with edges and points residual connections (EPResGCN), a more suitable network model for hemodynamic analysis, is constructed and the edge features of the graph structure are fully utilized to improve the predictive performance of the network. After training, EPResGCN achieves better results than other baseline networks on both the one-dimensional hepatic vessel dataset and the human aorta dataset based on centerline aggregation, which proves that the method in this paper is effective and can be simply and efficiently applied to other vascular structures.

## METHODS

An overview of the proposed method is shown in Fig. 1, and the overall framework has five main components: a) extracting the centreline model from the 3D blood vessel model and extracting the features corresponding to the centreline nodes, including the area of the cross-section in which the nodes are located, the topological relationships and distances between the nodes; b) measuring the velocity profiles of the blood vessel inlet and the boundary conditions of the outlet; c) using the vessel centreline model and boundary conditions to perform one-dimensional haemodynamic simulation to obtain the haemodynamic parameters; d) Construct the corresponding graph structure based on the vessel centreline model and embed the boundary conditions into the graph node features; e) The deep graph neural network EPResGCN model takes the constructed graph structure as the input and outputs the corresponding haemodynamic parameters at the graph nodes, with the haemodynamic parameters derived from the analysis based on finite element method as the target, and optimise the neural network using the loss function calculation method of multi-objective optimisation.

**Fig 1.**
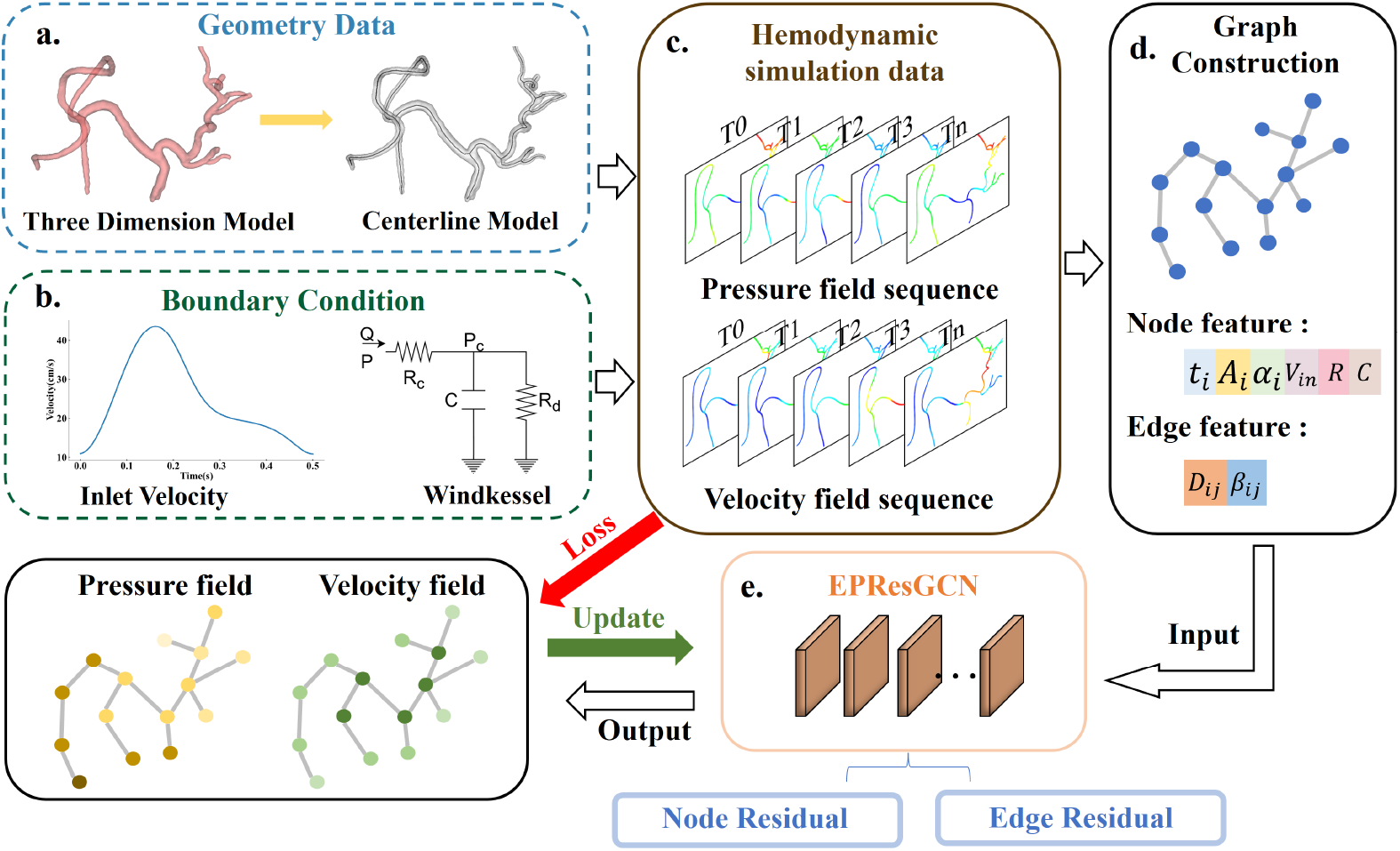
Overview of the pipeline used for one-dimensional prediction of hepatic vascular hemodynamics using EPResGCN. a: Hepatic vessel geometry data. b: Entrance velocity profile and boundary conditions. c: Hepatic vessel haemodynamic simulation data. d: Graph structure information. e: Deep neural network.

### Dataset

Data were obtained from Joint Innovation Laboratory for intelligent interventional procedures, Qingdao Municipal Hospital, all data have been approved by the clinical trial ethics Committee / institutional review board (IRB) (2024-KY-033), 24 sets of intraoperative hepatic CT images were collected, from which the vascular geometry model was extracted for modelling. Six of these sets of 3D vascular data are shown in Fig. 2 and the centre line is marked in it. The open source software 3D Slicer [36] was used to extract the vascular models from the CT images using the threshold segmentation method, and VMTK was later used to perform centreline extraction and feature calculation from the vascular models.

**Fig 2.**
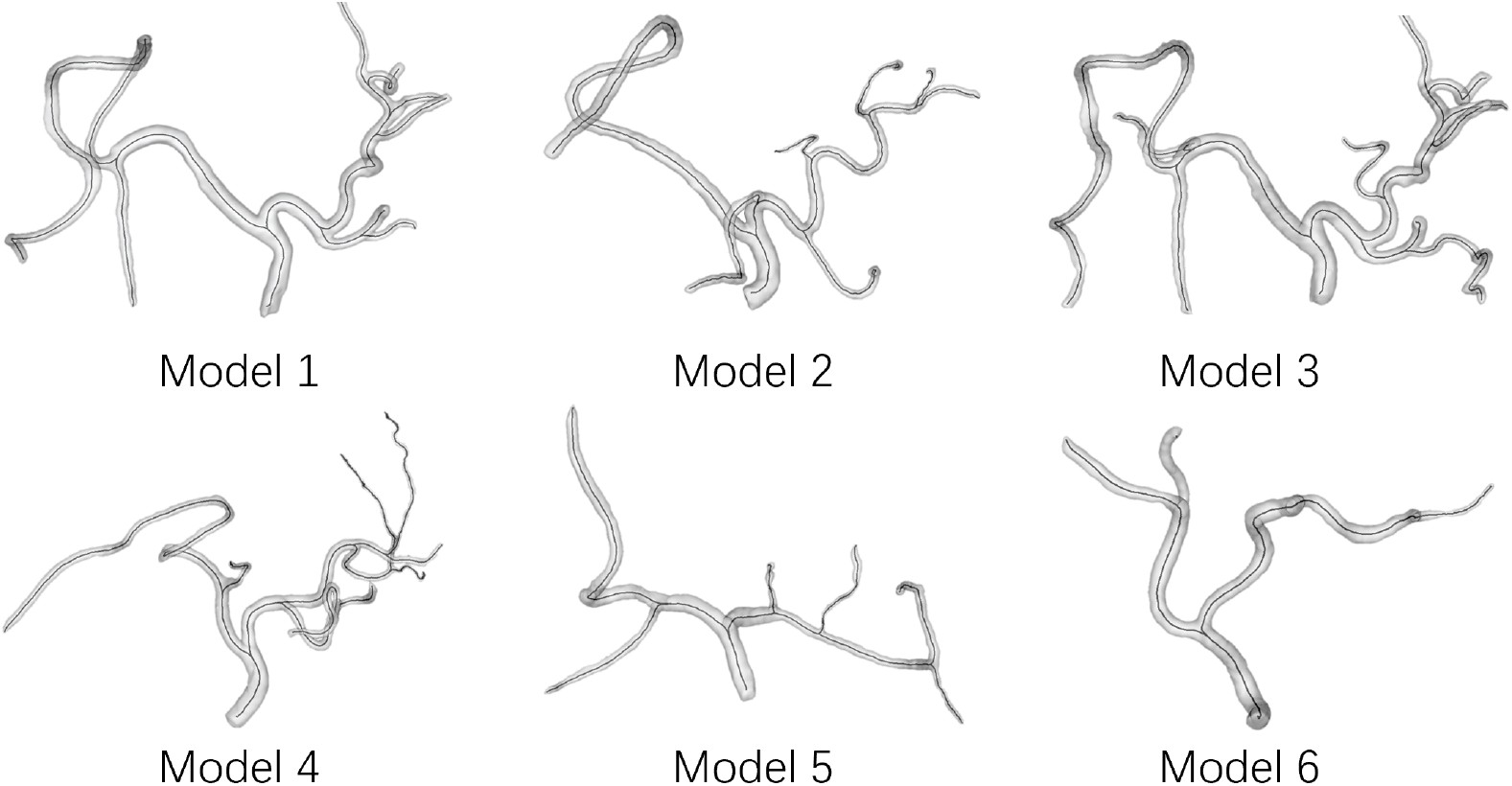
Geometric model of the hepatic vasculature. The curve in the vessel is the centreline.

Blood flow simulation of one-dimensional hepatic blood vessels was performed using Nektar++ [37] as a base tool, Nektar++ is an open source software framework that supports the development of a wide range of high-performance scalable partial differential solvers. In order to describe the velocities and pressures in the vascular network, it is assumed that the blood vessels are simplified as axially symmetric cylinders in the one-dimensional blood flow simulation, the blood is an incompressible fluid, and the vessel walls are deformable and impermeable pipes, and Eq. 1 is the controlling equation for the one-dimensional blood flow simulation.

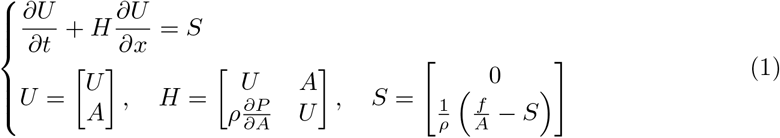

where *A* is the cross-sectional area of the vessel, *x* is the axial coordinate, *U* (*x, t*) denotes the axial velocity, *P* (*x, t*) denotes the intravascular pressure, *ρ* is the blood density, and *f* denotes the friction force per unit length. There are three unknowns in Eq. 1 as *U, A*, and *P*, so the pressure-area relationship equation is needed to close this system of equations:

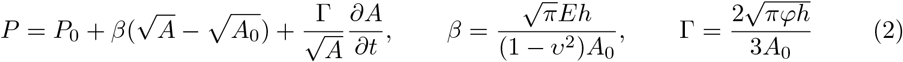

where *P*_0_ refers to the pressure outside the vessel, *A*_0_ is the initial cross-sectional area of the vessel, *E, h, v* and *φ* represent Young’s modulus, vessel wall thickness, Poisson’s ratio, and vessel wall viscosity.

The vascular network was modelled by obtaining the length of each vessel segment as well as the cross-sectional area of the origin from the hepatic vessel centreline model. The inlet boundary condition for the one-dimensional blood flow simulation was set as a pulsatile flow with a time period of 0.5, assuming conservation of mass and energy at the vessel bifurcation, and the outlet boundary condition was set as a three-element windkessel-type boundary condition [38]. Simulation calculations were performed using the DisContinuous Galerkin projection, the Runge-Kutta time integration method, and the Upwind format, with a simulation calculation time of 10 s. The selection of the The last 0.5 s of the steady calculation result is selected as the valid data. We performed 30 calculations with random fluctuations of 0.9-1.1 for inlet velocity and outlet boundary conditions for each set of vessels to construct our one-dimensional blood flow simulation dataset of hepatic vessels.

### Graph feature

Given a graph *G* constructed from the hepatic vascular centreline model, defined as *G* = (*X, E*), *X* ∈ ℝ^*N×F*^ is the node matrix, *N* is the number of nodes in the graph *G*, and *F* is the eigenvectors of the nodes. *E* ∈ ℝ^*N×N×P*^ is the edge matrix, the edge *E*_*ij*_ is the edge connecting two nodes *X*_*i*_, *X*_*j*_ between them, and *P* denotes the eigen-dimension of the edge between the two points. A set of graphs [*G*^0^, *G*^*t*^, …, *G*^*T*^] is used to represent a sequence of graphs over a cardiac cycle *T*.

#### Node feature

Based on the non-deformation of the translation or rotation of the physical system of the blood vessel, more attention is paid to the interrelationships and topological relationships between nodes when constructing the graph structure than to the coordinates of the nodes on the centreline, where the spatial location of the nodes will not be considered when constructing the node feature. *A*_*i*_ ∈ ℝ^+^ is the cross-sectional area of the section perpendicular to the centreline of the vessel, *i* is the node serial number. Each centroid on the centreline is classified as entrance node, outlet node, bifurcation node and branch node, one-hot coding *α*_*i*_ ∈ ℝ^4^ is used to represent the different node types. The boundary conditions 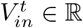 are the inlet velocity of the vessel at time *t*, with a three-element windkessel-type at the outlet, *R* ∈ ℝ is the sum of the vascular resistances *R*_1_ + *R*_2_ at the outlet points, and the value of *R*_1_ defaults to the characteristic impedance at the endpoints, *C* ∈ ℝ is the value of vascular compliance, and the values of *R* and *C* at the nodes inside the vessel are 0. The corresponding pressures 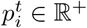 and velocities 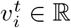 at the nodes are labels for the training process and are not used as inputs to the network. In summary, we obtain the node characteristics of the graph:

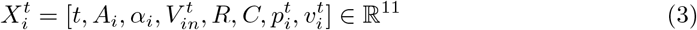

#### Edge feature

The distance between two points *D*_*ij*_ ∈ ℝ+ is the true distance between the two points along the centreline of the blood vessel, and a strategy of summing up the true distances of multiple edges is adopted for node downsampling. In this paper, we take the approach of embedding the boundary conditions into the boundary nodes, and in order to construct the connection between the internal points and the boundary points, in addition to the physical edges, the following artificial edges are added: the edges from the internal nodes to the outlet nodes, the edges from the internal nodes to the entrance nodes, and the edges from the branch start nodes to the inside of the branches. For each internal node, it is connected to its nearest outlet node or entry node at a distance *D*_*ij*_ which is the shortest path between the two points. The one-hot encoding *β*_*ij*_ ∈ ℝ^4^ is used to represent the four different edge types. In summary, we obtain the edge characteristics of the graph:

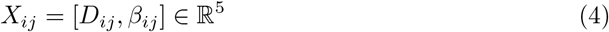

All edges are bidirectional, i.e. *E*_*ij*_ = *E*_*ji*_. The features other than *t, β*_*ij*_, and *α* are normalised, and the same features for all samples in the dataset are guaranteed to obey a Gaussian distribution *N* (0, 1). Note that *V*_*in*_ is in the normalisation process obey the distribution of velocity *V*, and the outlet boundary condition normalisation process only considers non-zero elements.

### Network architecture

#### Neural networks overview

The main body of the neural network consists of the encoding block, the network body EPResGCN, and the decoding block, using the superscript *l* to denote the output of the network at the *l*th layer, and the superscript *l*^*′*^ to denote intermediate results. The input node features and edge features are encoded using fully-connected neural networks (FCNNs): *X*_*en*_ = *F*_*pe*_(*X*) ∈ ℝ^*nk*^, *E*_*en*_ = *F*_*ee*_(*E*) ∈ ℝ^*mk*^, where *n* is the number of nodes, *m* is the number of edges, and *k* is the encoded feature dimension; the nodes and edges can have different encoded dimensions, and for the purpose of subsequent expression, the feature dimensions of both nodes and edges are mapped to *k*. The encoded *X*_*en*_ and *E*_*en*_ are used as inputs to the deep graph neural network, and after the multilayer graph convolutional neural network operation, the result is mapped to *X*_*de*_ ∈ ℝ^*n*2^ by the decoding block as the corresponding velocity and pressure fields.

#### Structure of EPResGCN

For graph neural networks, better results are not guaranteed by increasing the depth of the network; in fact, too much depth may lead to the appearance of the over-smoothing [39] phenomenon, which reduces the performance of the network and leads to worse results. Residual networks have been shown to be beneficial for training deep neural networks [40], we constructed a deep neural network EPResGCN that is more suitable for hemodynamic solution as shown in Fig. 3, with inputs of the network being the encoded node features *X*_*en*_ and the edge adjacency matrix *E*_*en*_.

**Fig 3.**
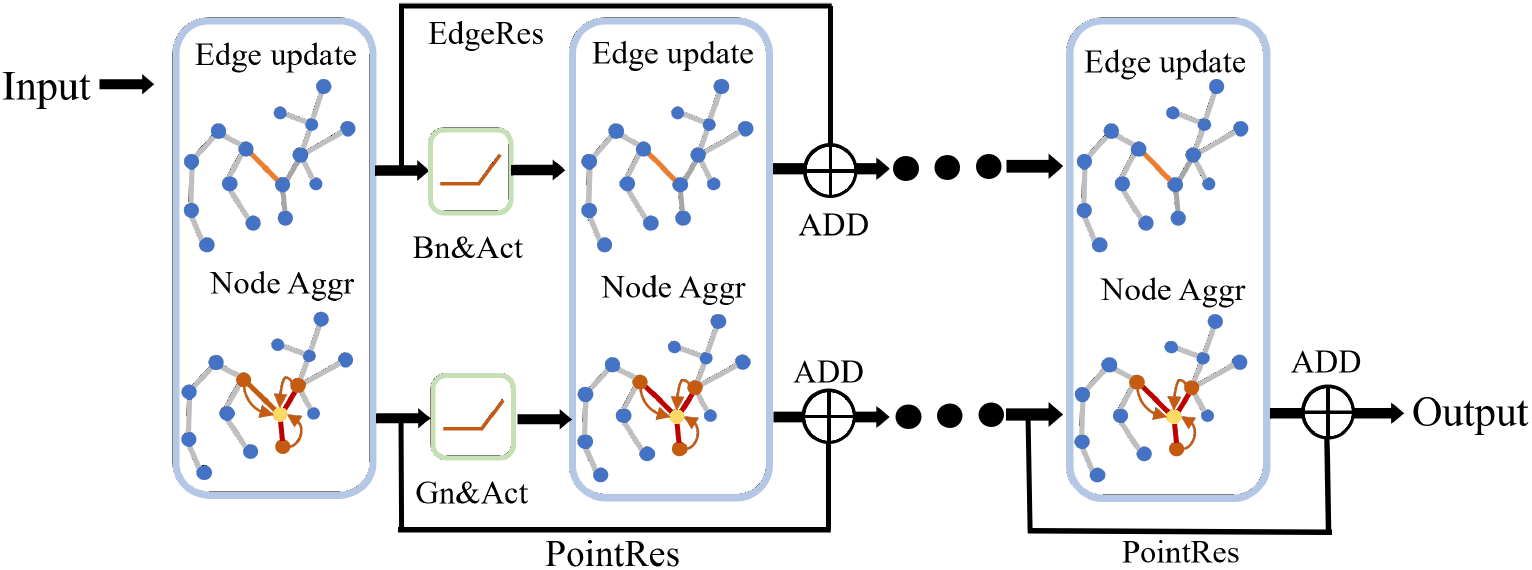
**The structure of EPResGCN**

Based on the importance of edge features in 1D haemodynamic analysis, if only node features are subject to network inference and residual connections, the edge features cannot be fully utilised, we propose to add node-like updating and residual connections to the edge features in the structure of deep graph neural networks to ensure that the utilisation of edge features is maximised. In constructing a deep neural network based on residual connectivity, we adopt the sequence of normalisation, activation, graph convolution, and residual connectivity. First, we compute new edge features:

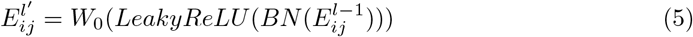

where *W*_0_ denotes the learnable weight matrix, *LeakyReLU* denotes the activation function, which has a small slope in the negative region to allow for some degree of negative transfer, and *BN* stands for batch normalisation. Due to the large amount of batch noise on the graph data, the batch statistics are not well concentrated around the dataset statistics, and the use of graph normalisation on the node features [41] allows the graph neural network to achieve faster convergence and better generalisation performance on the dataset.

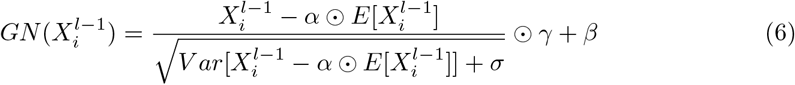

A learnable parameter *α* is used to automatically control the proportion of the mean that needs to be preserved, *σ* is a very small value to avoid a denominator of 0, *γ*, and *β* is an affine parameter, in the same way as for other normalisations.

Normalize and nonlinearly activate the node features of the previous layer to obtain a new node feature *X*^*l′*^ = *LeakyReLU* (*GN* (*X*^*l*^)), and aggregate the updated node features with edge features:

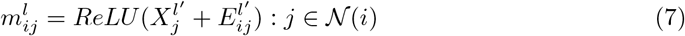

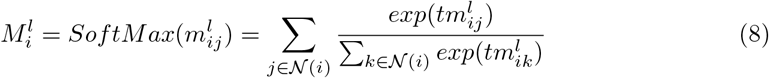

All the neighbour nodes 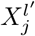 of the source node *i* and the corresponding edges 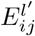 are summed to obtain the feature vectors 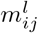, and all the feature vectors are aggregated by using the SoftMax aggregation function [42], where *t* is a continuous temperature coefficient variable. When *t* tends to 0 *SoftMax*(*t ·*) is equivalent to mean aggregation and when *t* tends to ∞ *SoftMax*(*t ·*) is equivalent to maximum aggregation.

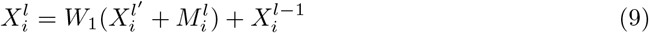

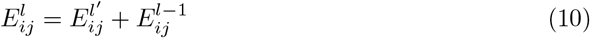

The feature vector of the source node itself is summed with the aggregated feature vectors of the neighbouring nodes, and the final output of this layer is obtained by residual connections of the linear output with the results of the previous layer, where *W*_1_ can consist of one or more weight matrices, and the edges are similarly joined with the edge features of the previous layer by a residual connections.

#### Loss function

For the pressures 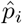 and velocities 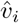 of the network’s outputs, MSE is used to compute their errors *p*_*loss*_ and *v*_*loss*_ with respect to the true labels *p*_*i*_ and *v*_*i*_, and weighted sums are employed to obtain the overall loss:

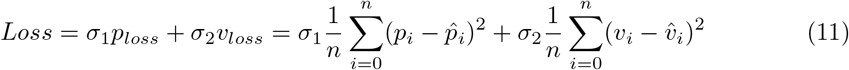

Defining the values of *σ*_1_ and *σ*_2_ using a manual approach is a huge challenge, and here we trade off multiple loss functions by considering the isotropic uncertainty of both [43].

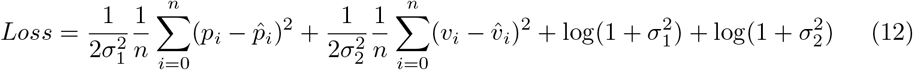

Define *σ* as a learnable hyperparameter that is adaptively tuned during training. To avoid mundane solutions, we enforce the regular term log 1 + *σ*^2^ to augment the *Loss*, where a large scale of *σ* will increase the contribution of the corresponding loss, and a small scale of *σ* will decrease the contribution of the corresponding loss, and no negative loss is incurred when *σ*^2^ *<* 1.

### Model evaluation and hyperparameter selection

We use the network’s predictions for the entire test set to introduce the error, which is computed using a symmetric mean absolute percentage error, where *m* is the number of samples in the test set and *n* is the number of nodes in a graph structure, and we compute the error for the test set at the end of each epoch.

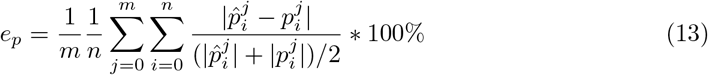

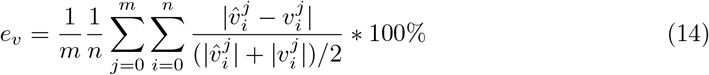

The choice of hyperparameters is crucial for the training of the neural network, the node and edge feature dimensions in both the coding and convolutional layers of the deep graph neural network are 256, the batchsize is set to 64, and the learning rate is annealed using an annealing algorithm, with an initial learning rate of 10^−3^ ending at 10^−5^, and 100 rounds of epochs for each training epoch.

## Result and analysis

In this paper, the prediction results of the networks are presented and analysed from four perspectives; in the first subsection, the results of haemodynamic prediction using different graph neural networks on the current dataset are explored in order to evaluate their effectiveness on the haemodynamic prediction task. In the second subsection, the effect of the depth of the graph neural network on the haemodynamic prediction results is investigated to explore the advantages of using edge residual connections in the haemodynamic prediction task. In the third subsection, the effect of using different connections in deep neural networks on network performance is tested. In the fourth subsection, haemodynamic parameters are predicted for each time step throughout the cardiac cycle using the trained graph neural network to investigate the performance of the network on long series of data.

### Comparison of different kinds of graph neural networks

In this section, we will demonstrate the performance of different graph neural networks in one-dimensional hepatic vascular hemodynamics prediction, including the depth based graph convolutional neural network EPResGCN, DeeperGCN [42], the attention based graph convolutional neural network Graph Transformer [44], GAT [45], GATv2 [46], and the non attention graph convolutional network CGdCNN [47], PDN [48], GraphGym [49]. All these network models were trained on the hepatic vascular dataset in this study, using the same baseline for training. For deep neural networks, we chose a model with a depth of 18 for training.

The prediction results of different graph neural networks on the same test data after training are shown in 4, in terms of error range, EPResGCN based on deep graph neural network performs the best (velocity error ±0.08, pressure error ±0.07), DeeperGCN is the second best (velocity error ±0.15, pressure error ±0.1), and the error map of the overall color is more uniform, and the pressure error and velocity error of the remaining networks range above ±0.2 and ±0.36, with more red and yellow areas. From the error plots, it can be obtained that the results of the deep graph neural network are significantly better than the shallow graph neural network, and the depth-based network performs better in terms of prediction accuracy and prediction stability.

After observation, it can be found that the prediction error is regional, which is expressed as a continuous larger error on the error graph. This is determined by the characteristics of the graph neural network, the graph neural network in the processing of graph structure data, usually for each node information transfer and aggregation operations, in this process, the prediction results of each node will be affected by its neighboring nodes, non-deep graph neural network due to the limited field of view, so when some nodes appear to have a larger error, it will affect the surrounding nodes, resulting in the entire region of the error are larger. The depth-based graph neural network, due to its large receptive field, for each point, the error of a few points within the receptive field has less impact on itself, so in the case of a relatively large error at an individual point, the impact on the neighbors is also smaller, as shown in the error map for the red region when the appearance of the range is small.

The numerical performance of the above graphical neural networks is shown in Table 1.Pressure error and velocity error are calculated by Equation 13 and Equation 14 respectively, and the overall loss error is the mean value of the pressure error and the velocity error.The graphical neural networks appearing in this section are trained using the same hyperparameters and data. The performance of the different networks in this task is evaluated by different evaluation metrics on the test set after the completion of training.The velocity error and pressure error of 1.593% and 1.521% for DeeperGCN and 1.317% and 1.293% for EPResGCN achieve better results among all the baseline networks. The prediction results of all networks pressure error is greater than velocity error because the input data uses inlet velocity and outlet boundary conditions which makes the network not as good at predicting pressure as velocity, in this regard deep neural network has a better result, the difference between pressure error and velocity error is at 0.072% for DeeperGCN and 0.024% for EPResGCN. According to the numerical results in the table, we can see that the depth-based network model can achieve a better result, which indicates that the deep network can better capture the complexity of hemodynamic parameters and provide more accurate prediction results, and for the addition of edge residual connections in the deep graph neural network can get a better result, which indicates that the edge residual connections contribute to the stability of the information transfer and the training of the model, thus improved prediction accuracy.

**Table 1.**
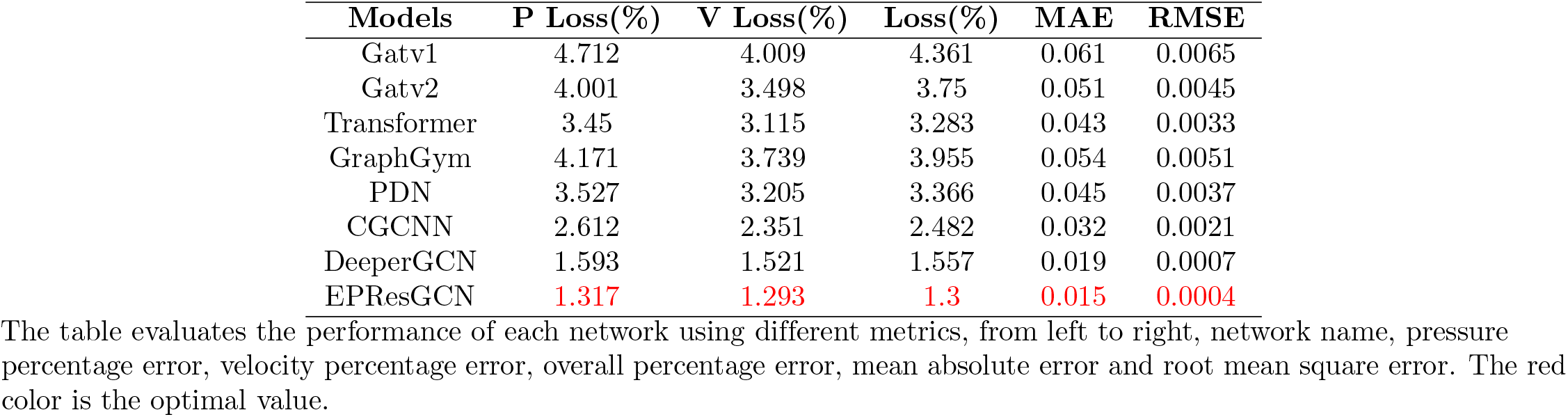
Tables of the performance of different networks.

It is worth noting that although the network based on the attention mechanism has a larger number of parameters, the difference between its results and those of the non-attention mechanism is very small, which is due to the special structure of the blood vessel network, the blood vessel from the entrance to the outlet of the blood vessel has a great deal of depth in the constructed graph structure, and except for the bifurcation point, all of them have only two neighbors: the upstream node and the downstream node, and even though we have added the artificial edges, each node still has very few neighbors, which leads to the attention mechanism not being fully utilized.

### Results of edge residual connections at different depths

In this subsection, the effect of the depth of the network on the performance of EPResGcn and PeeperGCN will be investigated, for both networks validated on the one-dimensional hepatic vascular blood flow dataset and on a reduced-order human aortic blood flow simulation dataset (based on cross-sectional averaging of three-dimensional blood flow simulation results), respectively. For all experiments, a ten-fold cross-validation method was used to evaluate the performance of the networks. During each training session, the same hyperparameter settings and data preprocessing methods were used, and the two network structures had the same number of parameters, except for EPResGcn which used residual connectivity on the edges. This ensures that different network models are compared under the same conditions and reduces the uncertainty of the experimental results. After completing the ten-fold cross-validation, the loss of each network model over all validation sets is calculated and the average of these evaluation metrics is computed as the final performance metric of the network model. To assess the reliability of the performance, this paper also shows 95% confidence intervals for these metrics to indicate the margin of error. Such an approach provides a more accurate representation of the predictive performance of the network model over the entire dataset and provides information on the confidence level of the performance estimates.

Figure 5 shows the performance comparison between EPResGCN and DeeperGCN with network layers of 4,7,11,18,25,40, i.e., on the test set, with the red curve and the green curve denoting the network error loss curves of EPResGCN and DeeperGCN, respectively. Observing the curves in the figure, it can be seen that the yellow curve is lower than the blue curve in both datasets, except that the accuracy of EPResGCN is slightly lower than that of DeeperCGN at a depth of 4. This indicates that adding edge residual connections in deep graph neural networks can improve the performance of prediction, and the addition of edge residual connections has achieved a better Results.

**Fig 4.**
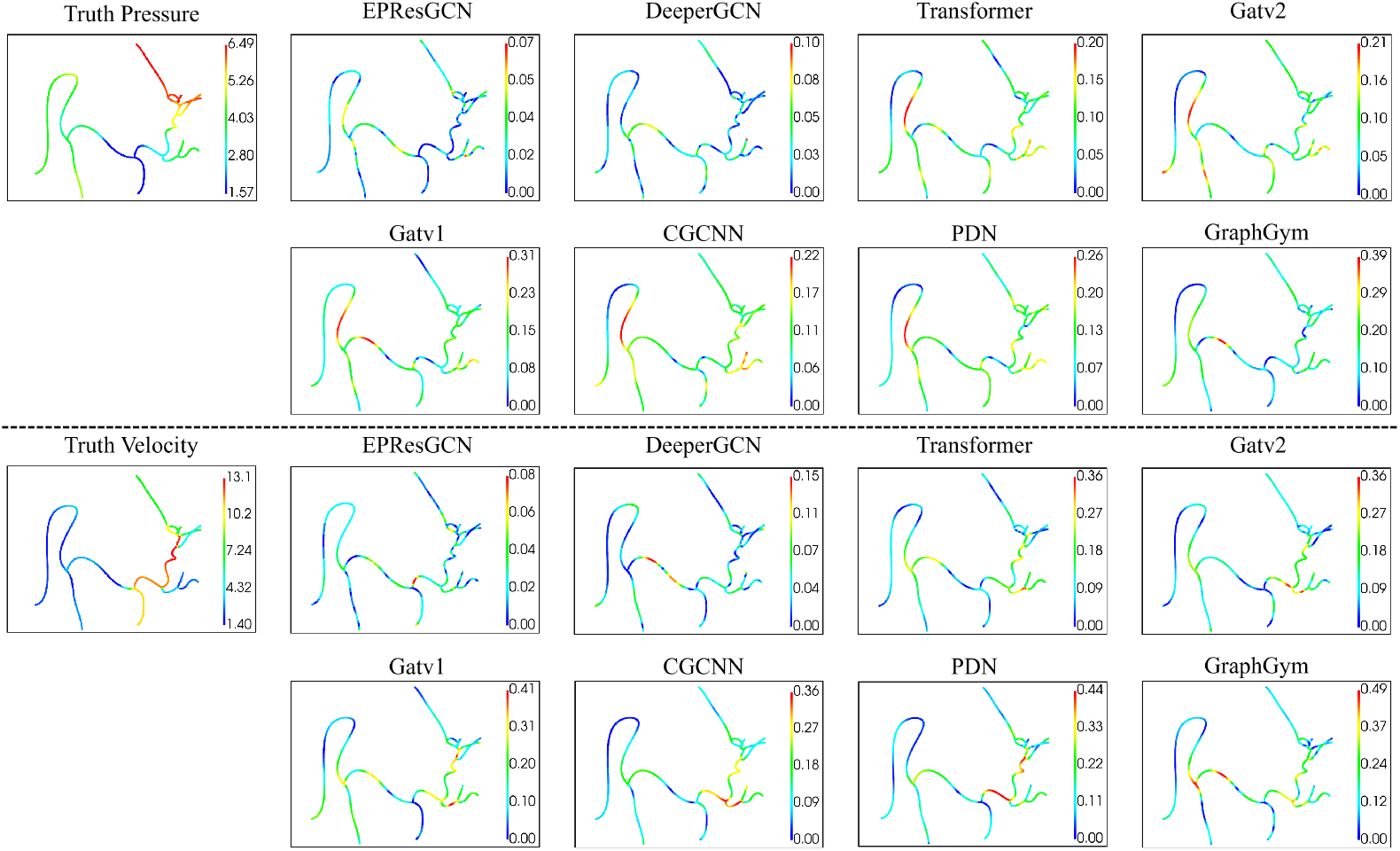
Distribution of errors in the prediction phase for different networks. The first column: the real values of velocity and pressure, columns 2-5: the error display of different network results, top: pressure error, bottom: velocity error. The left side of each cell shows the error distribution, the error size corresponds to the color blue to red (from small to large), and the value is the value corresponding to the colorbar on the right side, the unit of velocity is cm/s, and the unit of pressure is mmHg.

**Fig 5.**
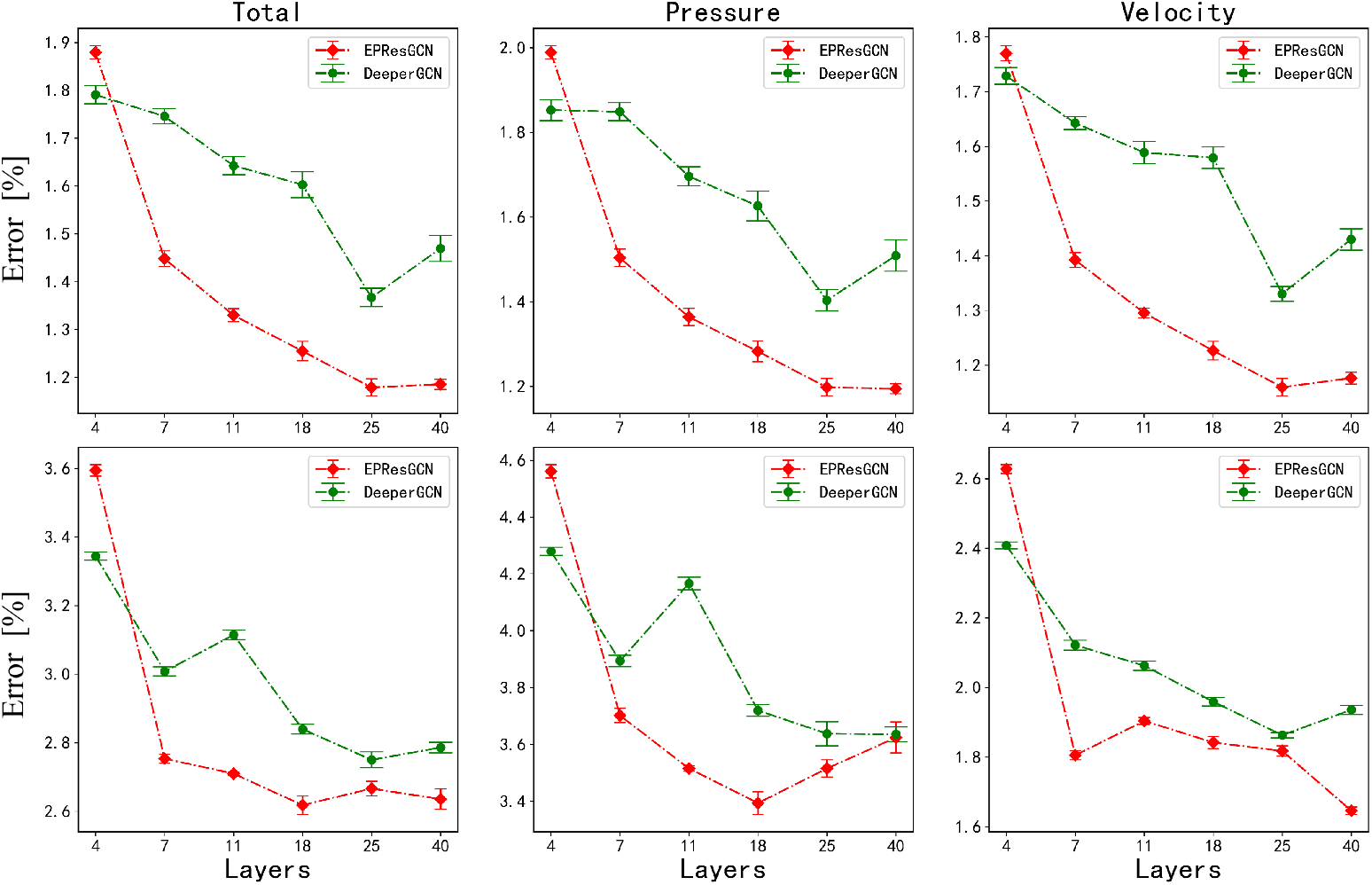
The effect of different neural network depths on the results. Upper: hepatic vessel dataset. Bottom: aortic dataset. Left: overall error. Center: pressure error. Right: velocity error. The red curve indicates the error of EPResGCN, the green curve indicates the error of DeepGCN, and the vertical line indicates the 95% confidence interval based on ten-fold cross-validation, where narrower confidence intervals indicate higher stability of the evaluation results, while wider confidence intervals may imply higher volatility of the performance.

The final results are improving with the increase of the number of network layers, which indicates that residual connections can improve the performance of the model by increasing the depth of the network and can effectively avoid the phenomenon of oversmoothing. However, with the increase in the number of layers, the performance improvement brought by the increase in computational effort is gradually reduced. Both EPResGCN and DeeperGCN reach a local optimum at 25 layers, and the results of DeeperGCN deteriorate after 25 layers, while the results of EPResGCN tend to converge, which suggests that EPResGCN is more advantageous in counteracting the oversmoothing phenomenon. By looking at the 95% confidence intervals of the prediction results on the hepatic vascularization dataset, it can be seen that the network with the added edge residual connections has smaller confidence intervals on the final prediction results, which indicates that the network has more stability.

A numerical representation of the experimental results in the hepatic vascular dataset is presented in Table 2, where the data are the mean values of the results of the corresponding ten-fold cross-corroboration, and the network with the addition of the edge residual connectings obtains an improvement of 0.43% from layers 4 to 7, exceeding the optimal results of DeeperGCN at layer 11, and as the depth of the network increases, the velocity error and the gap between the pressure error also narrows. Adding edge residual connections can improve the prediction accuracy and stability of the deep neural network while further narrowing the gap between velocity error and pressure error.

**Table 2.**
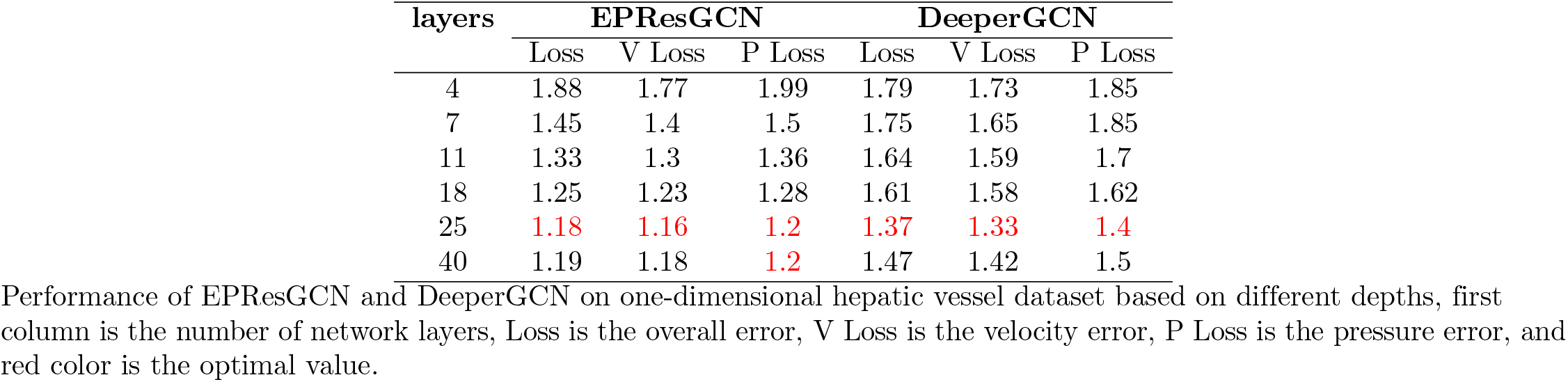
Results of EPResGCN and DeeperGCN at different depths.

| layers | EPResGCN |  |  | DeeperGCN |  |  |
| --- | --- | --- | --- | --- | --- | --- |
|  | Loss | V Loss | P Loss | Loss | V Loss | P Loss |
| 4 | 1.88 | 1.77 | 1.99 | 1.79 | 1.73 | 1.85 |
| 7 | 1.45 | 1.4 | 1.5 | 1.75 | 1.65 | 1.85 |
| 11 | 1.33 | 1.3 | 1.36 | 1.64 | 1.59 | 1.7 |
| 18 | 1.25 | 1.23 | 1.28 | 1.61 | 1.58 | 1.62 |
| 25 | **1.18** | **1.16** | **1.2** | **1.37** | **1.33** | **1.4** |
| 40 | 1.19 | 1.18 | **1.2** | 1.47 | 1.42 | 1.5 |

Performance of EPResGCN and DeeperGCN on one-dimensional hepatic vessel dataset based on different depths, first column is the number of network layers, Loss is the overall error, V Loss is the velocity error, P Loss is the pressure error, and red color is the optimal value.

### Effect of connection methods on deep graph neural networks

The experiments in this section are based on three network backbones to investigate the effect of different connection methods on the results, PlainGCN, DenseGCN, and ResGCN, the three network structures are given on the right side of Fig. 6. PlainGCN and ResGCN have the same number of network parameters except for the different connection methods, while DenseGCN has an expanded input dimension at each layer due to the splicing operation. DenseGCN has the same number of parameters as PlainGCN and ResGCN except for the different connection methods, and DenseGCN has the same number of parameters as PlainGCN and ResGCN due to the splicing operation which results in the expansion of the input dimensions of each layer.

**Fig 6.**
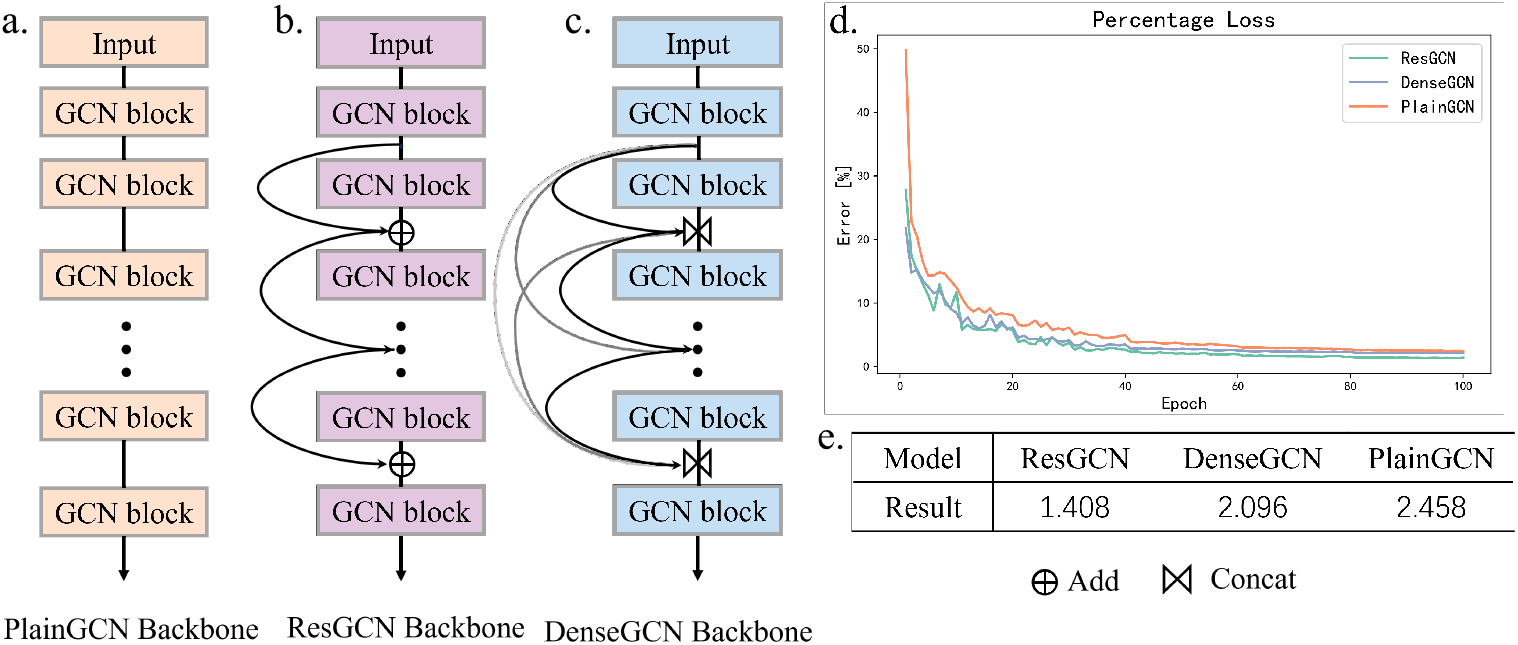
Network structure diagram and results. The network structure of PlainGCN. a: The network structure of ResGCN. b: The network structure of ResGCN. c: The network structure of DenseGCN. d: The error descent curves. e: The final results.

- **PlainGCN:** The network structure is a simple stacking of network layers, with no additional operations performed between layers.
- **ResGCN:**Residual concatenation is performed between the outputs of layers and layers, the dimensionality of node features and edge features are the same as PlainGCN, and the number of parameters of the network is the same as the number of parameters of PlainGCN.
- **DenseGCN:** By performing a dense connection between the outputs of each intermediate layer, the input of the *l* layer is spliced with the outputs of the previous *l* − 1 layer, and as the number of layers increases, the dimensionality of the input to the GCN module increases in equal proportions, where we control the dimensionality by setting an expansion parameter *D*, the dimension of the input to the *l* layer is *D*_0_ + *D* ∗ (*l* − 1).

Figure 6d gives a graph of the percentage error drop over 100 rounds for the graph neural network structures with the three types of connections. The green and blue curves are always lower than the yellow curve, indicating that both residual and dense connections help train deeper graph neural networks and effectively mitigate the over-smoothing phenomenon. The error values of ResGCN and DenseGCN are roughly the same before 30 rounds, and the errors of ResGCN based on residual connections are all smaller than those of DenseGCN based on dense connections after 30 rounds, and the final results are shown in the table in 6e. The results show that the residual connection is more effective than the dense connection on the one-dimensional hepatic vascular blood flow simulation dataset proposed in this paper.

### Prediction of the cardiac cycle

In this section, EPResGCN, DeepGCNs, Graph Transformer, Gatv2, CGCNN, and PDN are used to predict one cardiac cycle for six sets of blood vessels in graph 2. The cardiac cycle is set to 0.5s, with 0.02s as a time step, and a total of one hundred time steps, the trained graph neural network is applied to each time step within the cardiac cycle, and the performance of the network in predicting the cardiac cycle is evaluated based on the error between the network prediction and the labeling results.

Figure 7 shows the error curves of each network for the prediction of one cardiac cycle for six vascular models, calculating the velocity error and pressure error of the prediction results at each time step, and the curves represent the mean value of the error at each time step, which enables a more accurate assessment of the neural network model’s performance in the prediction of the cardiac cycle, by taking into account the error at each time step in the cardiac cycle. The mean curve reflects the overall prediction trend of the model, and it can be seen from the error plot that the depth-based graph neural network has the highest accuracy in predicting the cardiac cycle, EPResGCN’s velocity error and pressure error on the six vascular models are less than 1.5% except for the peak, and the peak error is also smaller, DeeperGCN’s overall error is less than 2%, but the peak error is larger. The results of CGCNN are slightly worse than DeeperGCN, but the fluctuation of the error is larger, with larger errors at multiple time steps, and the results of Graph Transformer, Gatv2 and PDN are worse, with error curves above 2% and larger peak errors. When predicting for Model 6, the pressure errors for all six models showed the largest errors in the early part of the cardiac cycle, with the largest errors exceeding 5% for EPResGCN and DeeperGCN, 15% for Gatv2, and 10% for the remaining networks.

**Fig 7.**
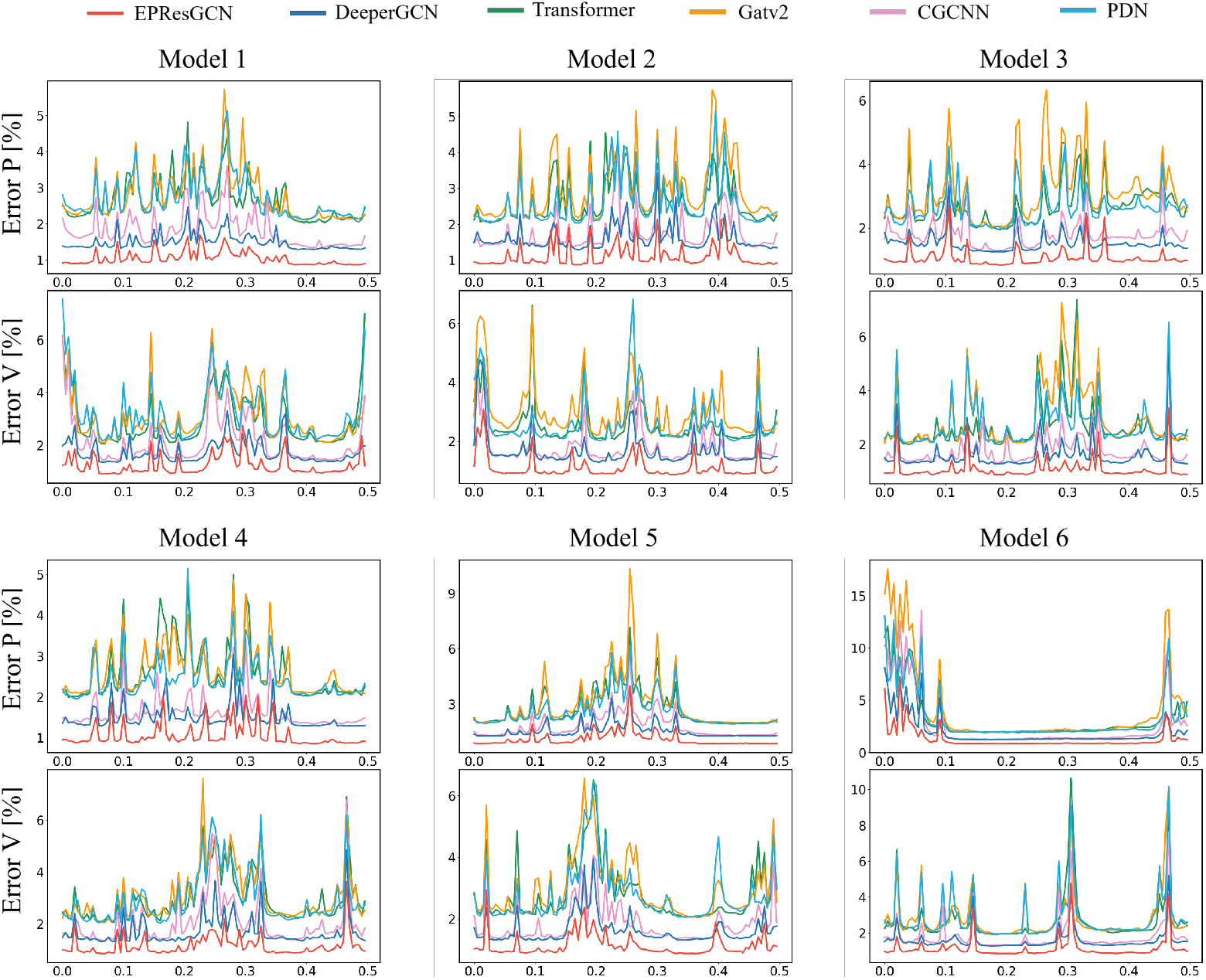
Prediction Error Plot for Cardiac Cycle. The horizontal coordinate is the time [s], the vertical coordinate is the error [%], and the solid line is the average of the errors at all nodes within each time step.

Error curves can be obtained EPResGCN performs best when predicting one whole cardiac cycle for each hepatic vascular model, the error curves maintain the lowest values relative to the other baseline networks, with less curve fluctuation. However, as with the other networks, the network model trained from independent time steps, due to its lack of temporal conditioning, suffers from an unsmooth error curve during prediction. This is due to the fact that at the beginning of the cardiac cycle, the blood flows into the blood vessels from the entrance of the heart, and the entrance velocity curve shows a low baseline level. With the contraction of the heart, the contractile force of the heart pushes the blood forward, and the velocity increases gradually, and the peak of the heart contraction is reached, and then it enters into the diastolic phase, and the blood flow rate slows down, and the entrance velocity curve of the whole cardiac cycle goes through a process of increasing and then decreasing. That is, in a cardiac cycle, there will be two time-step data with the same entrance velocity, although the node features contain time features, but the weight of the time features is less, resulting in the network for the two data with the same entrance velocity is sometimes difficult to differentiate, resulting in prediction error, which is manifested in the error curve is a sudden rise in the curve.EPResGCN has a wider sensing field, can better perceive changes in boundary conditions and time, and has the ability to differentiate between two time-steps of data with the same inlet velocity, so the error curve is relatively smoother.

## Conclusions

In this paper, we propose a neural network architecture for one-dimensional hemodynamic analysis of hepatic vessels. The neural network model is based on a deep graph neural network, which allows hemodynamic analysis of one-dimensional hepatic vascular structures. The dataset consists of hepatic blood vessels with different pathological features and there is a great geometric variability among the data samples. Experiments have demonstrated that the architecture proposed in this paper can also be applied to other one-dimensional vascular structures and has a strong generalization ability among different geometries. Comparing the three graph neural networks, depth-based, attention-based, and normal, respectively, the depth of the vascular map structure allows the deep graph neural network with a larger receptive field to achieve better results. We also found that because of the receptive field, the nodes of non-deep graph neural networks are more susceptible to the influence of their surrounding neighbors, which manifests itself as a larger error in regional continuity, which can be well overcome by deep graph neural networks. Subsequently, the edge residual connections were investigated, and in the comparison experiments of deep neural networks with different layers, the network models with added edge residual connections all achieved better results and more stable performance. Finally, using the same baseline trained graph neural network for the prediction of the whole cardiac cycle, the EPResGCN with the addition of edge residual connections and node residual connections achieves the best results in both single time step prediction and multiple time step prediction tasks within the cardiac cycle. Experiments show that the EPResGCN proposed in this paper can reduce both velocity error and pressure error to less than 1.5%, and for the prediction of the whole cardiac cycle, it can guarantee that the error of each time step is less than 2%.

While deep graph neural networks have better performance and show more stable performance in cardiac cycle multitemporal step prediction, a major disadvantage of deep neural networks is that they require more computational resources and computational time. The results in this paper show that the internal nodes are crucial for the perception of the boundary nodes, and the next step will be an in-depth exploration of the graph convolution kernel in order to construct a network model that is more suitable for the vascular structure, with the criterion of obtaining a larger sensory field while keeping the node’s own weights with a limited number of layers of the network, i.e., preventing the oversmoothing phenomenon due to the expansion of the sensory field. Embedding periodic boundary conditions into the graph structure and adding timing processing units to the network structure can better capture the dynamics and periodicity of time-series data, thus enhancing the model’s timing modeling capability, and based on the contextual information can improve the prediction capability of one of the time steps at each time step, resulting in a more accurate and stable prediction of the whole cardiac cycle. We will also enhance the richness of the hepatic vessel geometric model by modeling more patients’ geometric models and obtaining more realistic boundary conditions, so that more realistic situations can be taken into account in hemodynamic simulation, and richer data resources can make the network model have better geometric generalization performance.

## Supporting information

**S1 Fig. Overview of the pipeline used for one-dimensional prediction of hepatic vascular hemodynamics using EPResGCN**.a: Hepatic vessel geometry data. b: Entrance velocity profile and boundary conditions. c: Hepatic vessel haemodynamic simulation data. d: Graph structure information. e: Deep neural network.

**S2 Fig. Geometric model of the hepatic vasculature**.The curve in the vessel is the centreline.

**S3 Fig. The structure of EPResGCN**

**S4 Fig. Distribution of errors in the prediction phase for different networks**The first column: the real values of velocity and pressure, columns 2-5: the error display of different network results, top: pressure error, bottom: velocity error. The left side of each cell shows the error distribution, the error size corresponds to the color blue to red (from small to large), and the value is the value corresponding to the colorbar on the right side, the unit of velocity is mm/s, and the unit of pressure is mmHg.

**S5 Fig. The effect of different neural network depths on the results**.Upper: hepatic vessel dataset. Bottom: aortic dataset. Left: overall error. Center: pressure error. Right: velocity error. The red curve indicates the error of EPResGCN, the green curve indicates the error of DeepGCN, and the vertical line indicates the 95% confidence interval based on ten-fold cross-validation, where narrower confidence intervals indicate higher stability of the evaluation results, while wider confidence intervals may imply higher volatility of the performance.

**S6 Fig. Network structure diagram and results**.The network structure of PlainGCN. a: The network structure of ResGCN. b: The network structure of ResGCN. c: The network structure of DenseGCN. d: The error descent graph. e: The final results.

**S7 Fig. Prediction Error Plot for Cardiac Cycle**.The network structure of PlainGCN. a: The network structure of ResGCN. b: The network structure of ResGCN. c: The network structure of DenseGCN. d: The error descent graph. e: The final results.

**S1 Table. Tables of the performance of different networks**.The table evaluates the performance of each network using different metrics, from left to right, network name, pressure percentage error, velocity percentage error, overall percentage error, mean absolute error and root mean square error. The red color is the optimal value.

**S2 Table. Results of EPResGCN and DeeperGCN at different depths**.Performance of EPResGCN and DeeperGCN on one-dimensional hepatic vessel dataset based on different depths, first column is the number of network layers, Loss is the overall error, V Loss is the velocity error, P Loss is the pressure error, and red color is the optimal value.

